# Cortical sculpting of a rhythmic motor program

**DOI:** 10.1101/2025.06.20.660772

**Authors:** Eric A. Kirk, Kangjia Cai, Britton A. Sauerbrei

**Affiliations:** Case Western Reserve University School of Medicine, Department of Neurosciences

**Author notes:** These authors contributed equally (Eric A. Kirk, Kangjia Cai).

## Abstract

Motor cortex is the principal driver of discrete, voluntary movements like reaching. Correspondingly, current theories describe muscle activity as a function of cortical dynamics. Tasks like speech and locomotion, however, require the integration of voluntary commands with ongoing movements orchestrated by largely independent subcortical centers. In such cases, motor cortex must receive inputs representing the state of the environment and of subcortical networks, then transform these inputs into commands that modulate the rhythmic motor pattern. Here, we study this transformation in mice performing an obstacle traversal task, which combines a spinal locomotor pattern with voluntary adjustments that are impaired following cortical lesions. Cortical dynamics contain a prominent representation of motor preparation that is linked to obstacle proximity and robust to removal of somatosensory or visual input, and also maintain a representation of the state of the spinal pattern generator. These dynamics are driven by thalamic inputs through a low-dimensional communication subspace, which amplifies the preparatory transient. Cortical readout signals resembling muscle-like commands for obstacle traversal are small in amplitude, yet consistent across trials. Using computational modeling, we identify a simple algorithm that generates the appropriate commands through phase-dependent gating. Together, these results reveal a regime in which motor cortex does not fully specify muscle activity, but must sculpt an ongoing, spinally-generated program to flexibly control behavior in a complex and changing environment.

## 1. Introduction

To control voluntary movements, motor cortex must generate time-varying patterns of activity, then transmit commands to the spinal cord. For discrete movements such as reaching, cortex is the principal driver of motor output, and muscle activity can be described as a function of cortical firing rates (Fig. 1a, left)^1–4^. In many everyday tasks, however, cortical commands for voluntary actions must be integrated with commands generated independently by spinal or brainstem centers (Fig. 1a, right). For example, breathing is governed by a central pattern generator (CPG) in the medulla^5^, which rhythmically activates the respiratory muscles and is not usually under conscious direction. During speech, however, voluntary commands must reshape the activity of these muscles to accelerate the inspiratory phase and synchronize the breathing rhythm with syntactic breaks^6,7^. Similarly, in mammalian locomotion, a spinal CPG produces the basic motor rhythm, even in the absence of cortical input^8–10^. But when an animal must voluntarily modify its gait to step over an obstacle or traverse a horizontal ladder, motor cortex becomes essential for accurate control, and must flexibly generate commands to modulate the rigid and automatic pattern fixed by the CPG^11–17^.

**FIGURE 1.**
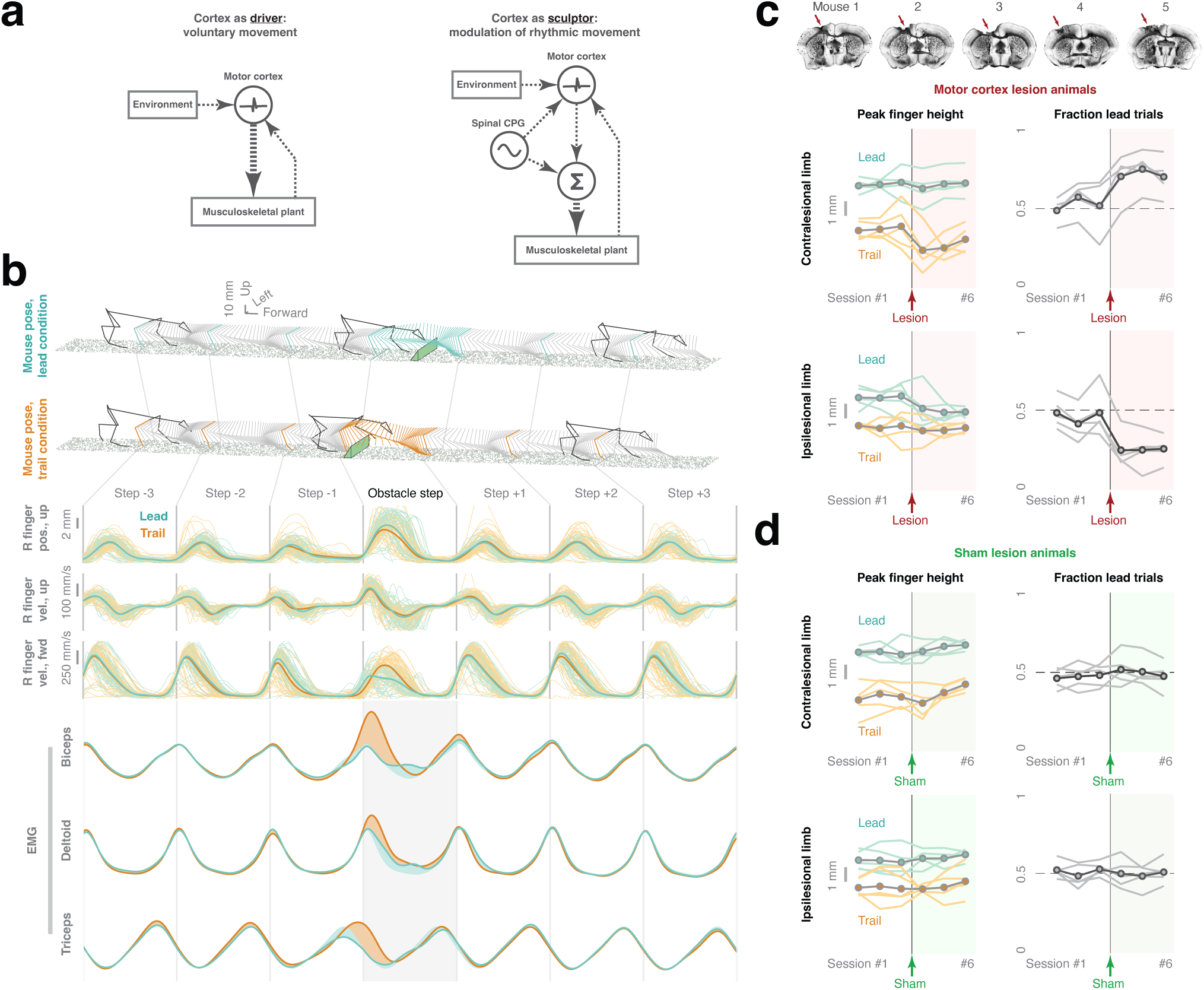
A cortically-dependent skilled locomotion task. **a**, Schematic illustrating a model of motor cortex as a driver of muscle activity (left), and an alternative model of cortex as a sculptor of a subcortically-generated rhythm (right). **b**, Limb kinematics and muscle activity in the task. Upper: three-dimensional pose estimates during obstacle traversals in the lead condition (blue) and trail condition (orange). Middle: position and velocity of the right finger aligned to step sequences in an example session. Lower: muscle activity of the biceps brachii, deltoid (anterior aspect), and triceps brachii across the sequence. Traces show the mean electromyogram (EMG) time series for each muscle in the lead and trail conditions (biceps, n = 8 sessions, n = 3 mice; deltoid, n = 4 sessions, n = 2 mice; triceps, n = 6 sessions, n = 3 mice). **c**, Effects of motor cortical lesions (n = 5 mice). Upper insert: histological sections showing lesions in each mouse. Left column: median peak finger height during obstacle traversal of the contralesional (upper panel), and ipsilesional (lower panel) forelimbs. Each curve shows a separate animal across three pre-lesion and three post-lesion sessions, and the dotted curve shows the average across animals. Right column: limb preference during obstacle traversal showing the fraction of trials in which the contralesional (upper panel) or ipsilesional (lower panel) limb led. **d**, Effects of sham lesions (n = 5 mice). Conventions are as in c.

Obstacle traversal clearly depends on cortical information processing^12,13,18–20^, but does not permit a description of muscle activity as a function of cortical firing rates, as the locomotor rhythm can continue without corticospinal drive. Motor cortex therefore acts not as a prime mover, but as a sculptor of a motor program produced independently by a subcortical pattern generator. How does the cortical network compute the appropriate commands for gait modification to reshape motor output? Here, we approach this question by combining large-scale neural recording in freely-moving mice, kinematic analysis, behavioral and optogenetic perturbations, and computational modeling. After verifying that obstacle traversal requires motor cortex using lesions, we identify population-level cortical dynamics related to the approaching obstacle, the locomotor rhythm, and the modulation of muscle activity during traversal. Next, we show that the cortical dynamics in advance of obstacle traversal represent motor preparation, rather than sensory input, while the rhythmic dynamics represent an efference copy from the CPG. We then demonstrate that signals resembling the preparatory and rhythmic factors are present in thalamic areas providing input to motor cortex, and identify a thalamocortical communication subspace which amplifies the preparatory transient. Finally, using a computational model, we specify a phase-dependent gating mechanism capable of translating the preparatory and rhythmic factors into commands that send the forelimb over the barrier.

## 2. Results

We trained freely-moving mice to perform a skilled locomotion task which integrates a spinally-driven, cortex-independent motor program with discrete, voluntary modifications of gait requiring motor cortex. Animals trotted on a linear treadmill, and were required to modify their limb and body motion to clear obstacles fixed to the treadmill belt. We used a high-speed multi-camera system and machine vision methods to estimate the three-dimensional pose of the mice and obstacles, and transformed the pose estimates into belt-centered coordinates in which mice locomote on an infinitely long linear track^21^. Because the belt speed was maintained at a constant value, these coordinates defined an inertial frame of reference. We then identified sequences of seven consecutive steps of the right forelimb in which the limb traversed the barrier on the central, fourth step (n = 5519 sequences, n = 22 sessions, n = 3 mice). Animals performed the task using two distinct strategies: initiating the step over the barrier with the right forelimb first, then following with the left forelimb (lead condition; Fig. 1b, top rows, blue), or leading with the left forelimb and trailing with the right (trail condition; Fig. 1b, top rows, orange). During obstacle traversal, peak fingertip height increased while forward velocity decreased, and these changes were larger in the lead condition (Fig. 1b, Extended Data Fig. 1a, top and middle, 1b). In the trail condition, animals shortened their stride one step in advance of the barrier (Extended Data Fig. 1a, lower), then briskly flexed the limb to increase upward velocity during traversal (Extended Data Fig. 1b). These kinematic modifications were generated by condition-specific changes in muscle activity, which were generally larger in the trail condition (Fig. 1b, lower, Extended Data Fig. 1c).

Is motor cortex necessary for performance of this task, and if so, what is its contribution? While several studies in cats and mice have documented deficits in obstacle traversal following lesions to motor cortex^13–16^, it is conceivable that the large number of trials and regularly-spaced obstacles in our version of the task resulted in consolidation of the control system to subcortical circuits^22,23^. Such consolidation has been observed in rodents trained to produce brief but precisely-timed and highly stereotypical movement sequences over tens of thousands of trials^24–26^. Our kinematic data suggested the animals were not memorizing a precise motor sequence and repeating it in open loop: animals frequently stopped and started locomotion between obstacles, lead and trail sequences appeared to be interleaved randomly, and the number of steps between consecutive obstacles varied across trials (Extended Data Fig. 1d). Nevertheless, we directly tested the contribution of motor cortex by assessing performance before and after an aspiration lesion to this area (see Methods). Following a lesion, animals (n = 5 mice) exhibited a robust drop in peak fingertip height over the obstacle with the contralesional limb when this limb trailed (Fig. 1c, left column, upper; p = 4e-3, rank sum test), and a small drop in height for the ipsilesional limb when this limb led (Fig. 1c, left column, lower; p = 3e-3, rank sum test). Animals also began to prefer leading with the contralesional limb and (by symmetry) trailing with the ipsilesional limb (Fig. 1c, right column; p = 4e-4 and 6e-5, respectively, rank sum test). None of these effects were observed in a group of control animals that received sham surgeries which included a craniotomy but no lesion (Fig. 1d, n = 5 mice; all p-values > .05). These data demonstrate that execution of the obstacle traversal task requires motor cortex. Furthermore, the strong asymmetry of effects on the ipsilesional and contralesional limbs, along with the larger kinematic impairment in the contralesional trail condition, which requires larger changes in muscle activity (compare Fig. 1c, upper right, to Fig. 1b, lower), suggests the role of motor cortex in this task is the generation of descending commands which activate specific muscle groups.

Execution of this skilled locomotion task requires the motor cortex to perform a key computation. It must receive a transient input conveying the proximity of the obstacle and a rhythmic input encoding the phase of the locomotor rhythm, then transform these inputs into an output carrying descending commands that modify muscle activity and propel the limb over the barrier. Neither of these inputs alone is sufficient to produce an appropriate motor command. If the rhythmic input were preserved while the transient input were suppressed, the cortex would not be informed of the need to clear the obstacle. Conversely, if the transient input immediately triggered movement irrespective of locomotor phase, the animal would stumble.

How does motor cortex implement this computation? To address this question, we performed large-scale recordings in freely-moving animals by chronically implanting 5120-contact, 384-channel Neuropixels 2.0 probes^27^ in forelimb sensorimotor cortex^28–30^ of the left hemisphere, contralateral to the reference limb (Fig. 2a). This approach enabled us to record from large ensembles of neurons simultaneously (n = 5596 neurons, n = 22 sessions, n = 3 mice). Cells exhibited a wide range of temporally-precise, task-locked responses. Some fired rhythmically during locomotion on a flat surface, but were minimally influenced by obstacle traversal (Fig. 2b, neuron 7), while others had transient responses. In some cases, the amplitude of the transients in the lead and trail conditions differed (neurons 4 and 5), while in others it was similar (neurons 6 and 8). Furthermore, the timing of the transient with respect to the phase of the contralateral limb was in some cases preserved between lead and trail conditions (i.e., was limb-dependent; neurons 2 and 8), and in others aligned with swing onset for the first limb to traverse the barrier, independently of whether the limb was contralateral or ipsilateral to the neuron (Fig. 2b, neuron 6). Many cells also exhibited superpositions of rhythmic and transient responses (Fig. 2b, neurons 1, 2, and 3). To further quantify the responses of single neurons, we classified the cells using three criteria (Fig. 2c, Extended Data Fig. 2; see Methods): (1) the effect of step number within the sequence on firing rate (67% of cells significantly modulated; two-way ANOVA with Benjamini-Hochberg correction for multiple comparisons, q < .05), (2) the effect of an interaction between step number and lead vs trail condition on firing rate (21%; two-way ANOVA, q < .05), and (3) synchronization of spike times with the locomotor rhythm for steps on a flat surface (59%; Rayleigh test, q < .05). A large majority of neurons (83%) met at least one criterion, and many met two (36%) or three (14%) criteria. This range of responses across individual neurons is broadly consistent with findings in the cat primary motor^12,13^ and premotor^18,31,32^ areas, which contain cells rhythmically active on a flat surface, modulated when the trajectory of the contralateral limb must be altered, and transiently active in advance of obstacle clearance.

**FIGURE 2.**
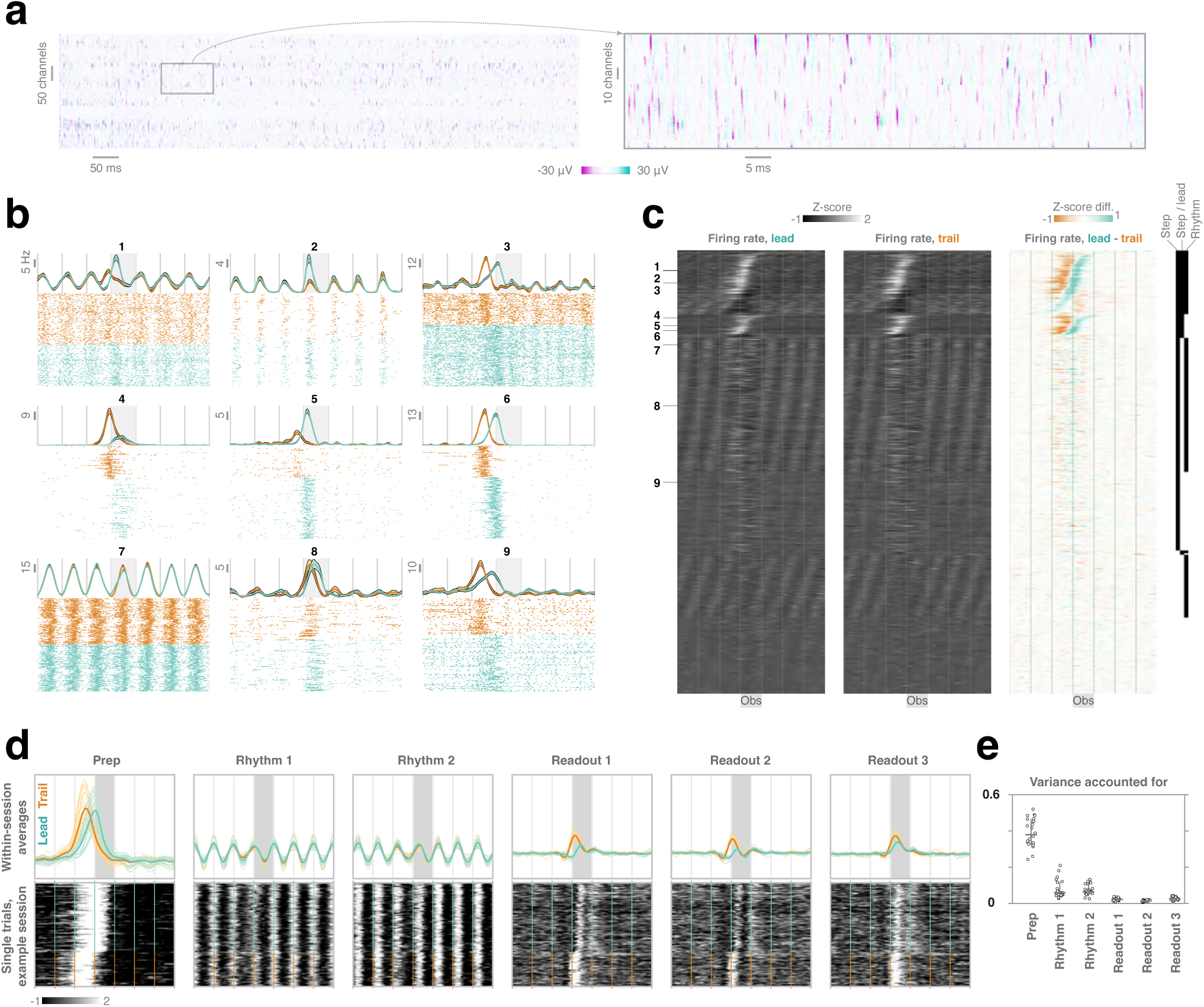
Task-relevant dynamics in motor cortex. **a**, Raw electrophysiological data recorded with Neuropixels 2.0 probes chronically implanted in forelimb motor cortex of the left hemisphere. Rows correspond to channels, columns to time, and color to voltage. Data windows are 1 s (left) and 100 ms (right). **b**, Firing rates and raster plots for representative motor cortical units in the task. Firing rate error bars denote standard error of the mean. **c**, Responses of all single units in the task (n = 5596 neurons, n = 5519 sequences, n = 22 sessions, n = 3 mice). The left two panels show firing rate Z-scores as a function of step phase across the sequence of seven steps in the contralateral lead and trail conditions. The right panel shows the difference in firing rate Z-score between the lead and trail conditions. The inset at the far right indicates which neurons show firing rate modulation by step number in the sequence, by the interaction between step number and lead / trail condition, and by the locomotor rhythm. **d**, Population activity along factors related to motor preparation, the locomotor rhythm, and muscle-like readout. Upper: trial-averaged activity. Light lines correspond to single sessions (n = 22 sessions, n = 3 mice), and bold lines are the grand mean across sessions. Lower: single-trial dynamics within a single session. **e**, Fraction of total firing rate variance accounted for by each of the factors. Each point corresponds to a single session. Lines indicate median and interquartile range across sessions.

The activity of single neurons was temporally complex and highly diverse, consistent with findings in other tasks and species^33^. We therefore sought to identify neural factors - linear combinations of the firing rates of all simultaneously-recorded neurons - which express coherent patterns of activity across the population in coordinates interpretable in terms of task variables^34–36^. We first approached this problem using principal component (PC) analysis, which provides orthogonal coordinates capturing the maximum possible variance in the original firing rates. The top 15 PCs accounted for 90% of the firing rate variance (median across n = 22 sessions; Extended Data Fig. 3a). In every animal and session, the first PC exhibited a large, limb-independent transient in advance of the obstacle. In the lead condition, the peak in this transient coincided with swing onset for the contralateral forelimb; on trail sequences, it occurred half a step cycle earlier, when the ipsilateral limb initiated swing over the obstacle (Extended Data Fig. 3b-c). In most sessions, the second PC consisted of a biphasic, limb-independent transient, while PCs 3-6 were typically sinusoidal on all steps across the sequence. Higher PCs contained primarily smaller, transient signals, including limb-dependent transients at obstacle clearance (e.g., PC 9, Extended Data Fig. 3b).

Although principal component analysis was modestly effective at separating rhythmic and transient features of neural population activity, these features were frequently intermingled in the higher components, and sometimes varied between animals. We therefore used a targeted dimensionality reduction approach to isolate three types of consistent and interpretable population-level factors: a preparatory factor activated just before the animal traversed the barrier, rhythmic factors representing the cosine and sine of locomotor phase throughout the sequence, and readout factors activated when the contralateral limb initiates stepping over the obstacle (Fig. 2d, Supplemental Video 1; see Methods). The preparatory factor, defined as the first principal component score, accounted for 38% of the total firing rate variance (median across n = 22 sessions; Fig. 2e). This factor was activated before the onset of the step over the barrier, independently of whether this step was initiated with the contralateral or ipsilateral limb. Thus, we conclude the preparatory factor is not a motor command for modulating activity in the contralateral forelimb muscles. The two rhythmic factors exhibited sinusoidal patterns^12,13,19,37^ which were not modulated by obstacle traversal (Fig. 2d, top row), and accounted for 6% and 7% of the firing rate variance, respectively (medians across sessions). While these signals resemble aspects of limb kinematics, they are also unlikely to constitute motor commands, as locomotor performance on a flat surface is essentially unchanged by lesions or inactivation of motor cortex^13–16,38^. To identify signals which could instruct the cortically-dependent changes in muscle activity during obstacle traversal, we therefore extracted cortical factors representing deviations in EMG from the stereotyped, rhythmic pattern (Extended Data Fig. 1c; see Methods). These readout factors differed in amplitude between lead and trail conditions at contralateral swing onset (Fig. 2d), accounted for only 1-3% of the total firing rate variance (Fig. 2e), and were consistent across trials (Fig. 2d, lower) and sessions (Fig. 2d-e). The relative magnitude of the dominant factors (i.e., preparatory and rhythmic) and the smaller readout factors is reminiscent of observations in nonhuman primates performing a rhythmic cycling task^39,40^, where the largest signals in motor cortex do not resemble muscle activity, while the readout factors used to decode motor output account for only 3% of the variance.

Does the timing of the preparatory factor coincide with the proximity of the obstacle, or the initiation of the leading limb’s step over the barrier? To distinguish between these possibilities, we examined the relationship between preparatory activity and kinematics on single trials. During traversals on which the animal’s nose was more advanced relative to the obstacle at the onset of swing in the contralateral limb, the preparatory transient occurred earlier in the step sequence (Fig. 3a-b; p = 4e-5, signed rank test against null hypothesis of zero median correlation). This transient was therefore synchronized more tightly with obstacle proximity than with motion of the leading limb. Furthermore, when the animal failed to smoothly execute a sequence of seven steps, but paused at the obstacle, the preparatory factor remained elevated for the duration of the pause (Fig. 3c-d; p = 5e-5, signed rank test against null hypothesis of zero median correlation). These observations are consistent with the hypothesis that the neural factor represents an abstract motor intention. Alternatively, the factor could reflect a relatively direct propagation of sensory information from the whiskers or from vision. Preparatory dynamics were maintained during optogenetic silencing of barrel cortex, which processes sensory input from the whiskers and communicates with the forelimb motor area (Fig. 3e-f). The possibility remained, however, that motor cortex could be driven by vibrissal sensory signals from subcortical areas, as suggested by prior behavioral and lesion experiments^16^. To test this hypothesis, we recorded activity from the same neurons as the animal performed the task with intact whiskers and following contralateral and then bilateral whisker trimming. Preparatory dynamics were robust to this removal of sensory feedback (Fig. 3g). Finally, these dynamics were maintained when the animal performed the task in the dark with intact whiskers (Fig. 3h). These findings demonstrate that the preparatory factor does not encode modality-specific sensory information, but is an abstract representation of motor intention.

**FIGURE 3.**
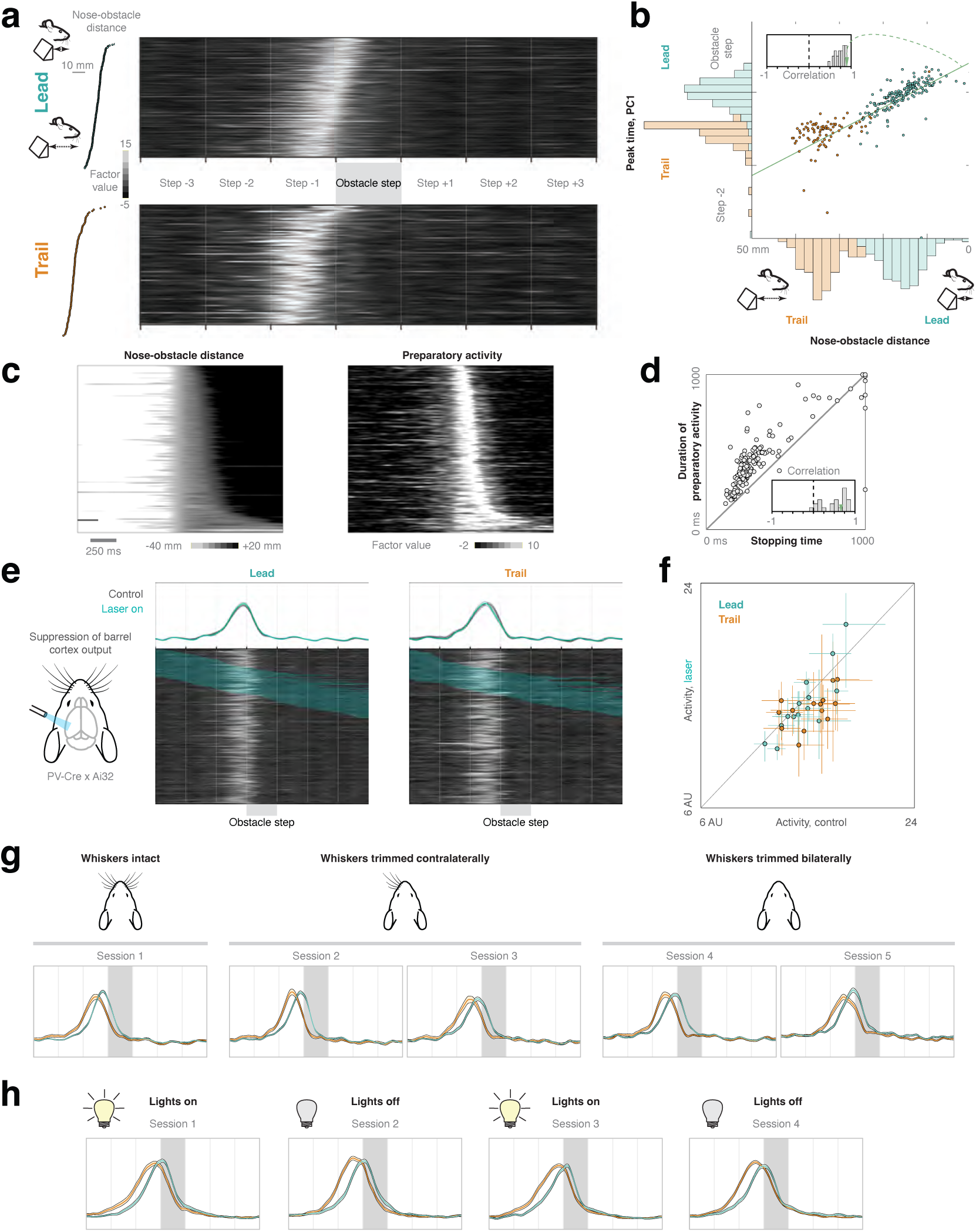
Preparatory dynamics are triggered by obstacle proximity and robust to sensory deprivation. **a**, Preparatory factor activation in a representative session, with trials sorted by the distance between the nose and the obstacle at swing onset for the contralateral forelimb. Insets display the nose-obstacle distances on single trials. **b**, Correlation between the timing of the peak in preparatory activity and the nose-obstacle distance. Inset shows the distribution of rank correlation coefficients across all sessions (n = 22 sessions, n = 3 mice). **c**, Left: nose-obstacle distance as a function of time, centered on obstacle crossing. Rows correspond to single trials, and are sorted by the duration for which the animal pauses at the obstacle. Right: Activation of the preparatory factor on the same trials. **d**, Correlation between the duration of the animal’s pause at the obstacle and the duration of the preparatory signal. Scatterplot shows all trials from a single session, and inset shows the distribution of rank correlation coefficients across sessions. **e**, Preparatory activity is robust to optogenetic inactivation of barrel cortex (n = 5543 neurons, n = 4442 sequences, n = 14 sessions, n = 1 mouse). Heatmaps display the factor values on single trials. Blue shading indicates laser-on epochs. Trials are sorted by the timing of the optogenetic stimulation. Upper panels display the mean preparatory activity on control and inactivation trials. **f**, Preparatory activity on inactivation vs. control trials for all sessions. Values represent the median, across trials, of the norm of PC 1 on the third and fourth steps. Error bars denote bootstrapped 95% confidence intervals. **g**, Preparatory activity is robust to whisker trimming. Panels show the mean preparatory factor across individual sessions using the same principal component coordinates (n = 250 neurons linked across five sessions, n = 1804 sequences, n = 1 mouse). Whiskers were trimmed contralaterally after the first session, and bilaterally after the third. Error bars indicate standard error of the mean. **h**, Preparatory activity persists in darkness (n = 241 neurons linked across four sessions, n = 1028 sequences, n = 1 mouse). Conventions as in g.

We next investigated the origin of the prominent signals entrained to the locomotor rhythm across the entire sequence of steps (columns 2-3 in Fig. 2d). In particular, we asked whether the cortical rhythm is driven by sensory feedback (e.g., from muscle stretch receptors), or instead by a copy of internally-generated commands. The sensory feedback hypothesis predicts that timing relationships between pairs of neurons during passive sensory stimulation should be preserved when the animal engages in voluntary movement. For example, if two neurons are activated by stretch in elbow extensor muscles when the animal is resting and the limbs are moved passively by the experimenter, they should also be co-activated at the same phase of the locomotor cycle during active movement. On the other hand, large changes in correlations between neurons across active and passive limb movement would suggest that the cortical rhythm during locomotion is at least in part internally-generated. We directly tested these hypotheses by recording from the same motor cortical neurons during the treadmill locomotion task and during passive sensory stimulation, in which the experimenter repeatedly cycled the forelimb in a pattern mimicking locomotion as the animal rested (Fig. 4a; see Methods). While the strength of cells’ entrainment to active and passive movement was correlated (Fig. 4b, upper; Spearman’s ρ = 0.55, p < 1e-10), we observed no relationship between the preferred movement phase between tasks (Fig. 4b, lower, Fig. 4c; circular-circular correlation = −0.02, p = 0.69). Similarly, while strong pairwise correlations between neurons were present in both active and passive movement, we observed a reorganization of the correlation structure between tasks (Fig. 4d). At the population level, we identified subspaces related to rhythmic active movement (as described above; c.f. Fig. 2d) and to passive movement by linearly decoding the cosine and sine of movement phase from firing rates. During locomotion, neural dynamics were confined to the active subspace, but remained constant across the movement cycle in the passive subspace. Conversely, activity during passive cycling was prominent in the passive subspace, but revealed a minimal footprint in the active subspace (Fig. 4e-f). Taken together, these results suggest that rhythmic cortical activity during locomotion primarily reflects internally-generated signals, rather than sensory feedback from the limbs^41^.

**FIGURE 4.**
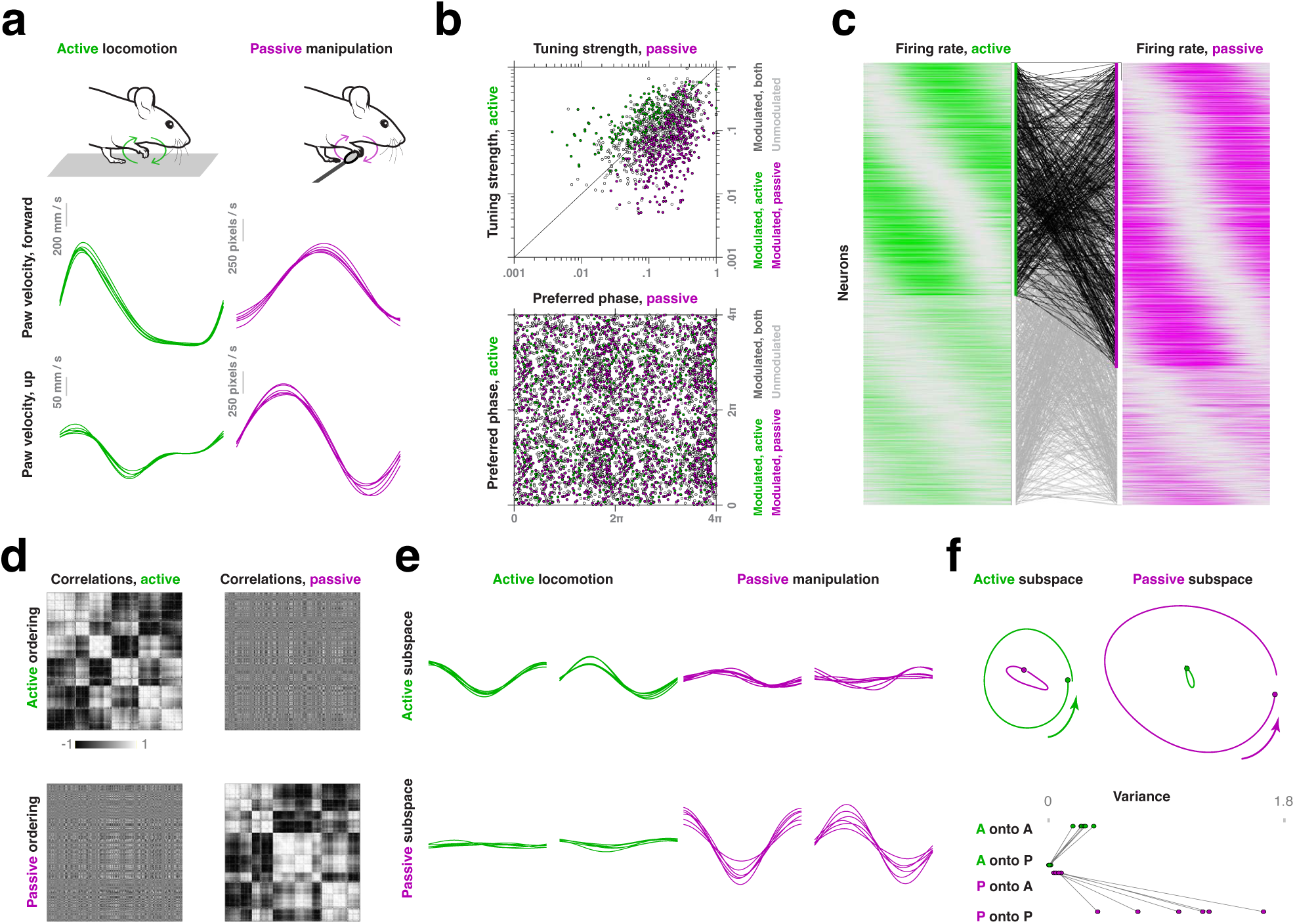
Central origin of the motor cortical rhythm. **a**, Finger velocity during locomotion on a flat surface (green, left) and during passive limb cycling (magenta, right). Each trace corresponds to the average of all trials in one session. **b**, Upper: degree of synchronization of spiking (mean resultant length) to the locomotor rhythm vs. to passive cycling. Each point corresponds to one neuron. Colors indicate which units are significantly modulated by locomotion, passive cycling, both, or neither. Lower: phase preference (mean resultant angle) for all units in active locomotion vs. passive cycling. **c**, Normalized firing rates for all units during active locomotion (green, left) and passive cycling (magenta, right). Bold lines indicate which cells are significantly modulated in each task. Rows are sorted first according to the detection of significant rhythmicity, then by preferred phase. Lines show the same neurons across the two tasks (n = 1263 neurons linked across active and passive conditions, n = 1973 sequences, n = 9865 steps across a flat surface, n = 2906 passive limb cycles, n = 6 sessions, n = 1 mouse). **d**, Firing rate correlations for all pairs of neurons. The top row displays correlations during locomotion, and the bottom row correlations during passive cycling. Neurons are sorted based on a hierarchical clustering of correlations during locomotion in the left column, and during cycling in the right column. **e**, Projection of firing rates during active locomotion and passive cycling onto the active and passive subspaces. Each trace corresponds to data from one experimental session. **f**, Upper: neural trajectories in the active and passive subspaces during the two tasks. Neurons and trials are pooled across sessions. Lower: variance of neural activity in each task and subspace.

So far, we have assumed that the transient and rhythmic dynamics are not generated de novo in motor cortex, but are driven by external inputs. What brain areas provide these inputs, and how are the dynamics in the communication channel^42–44^ related to those in the input regions and motor cortex? To address these questions, we implanted animals (n = 4 mice) with Neuropixels 2.0 probes targeted to two major thalamic inputs to motor cortex: the ventral anterior lateral complex (VAL; n = 215 neurons), which receives cerebellar and basal ganglia inputs, and the posterior complex (PO; n = 540 neurons), a higher-order nucleus involved in sensorimotor processing. Neuron locations were verified by registering histological sections to the Allen Mouse Brain Common Coordinate Framework, and by examining the depth profile of electrophysiological data across the probe shanks (Fig. 5a-c, Extended Data Fig. 4a-d; see Methods; Supplemental Video 2). The activity of single neurons in VAL and PO was strongly modulated by locomotor phase (87% and 85%), consistent with prior findings in the cat^45,46^, and by the interaction between step number and lead vs trail condition (79% and 82%; Fig. 5d-e, Extended Data Fig. 5a-b). Firing patterns in VAL and PO were highly similar, so cells were pooled for subsequent analysis. At the population level, the dominant factors identified with principal component (PC) analysis were near-sinusoidal (PCs 1-2, 20% and 14% of variance), and were followed by smaller, transient signals synchronized with obstacle proximity (PCs 3-4, 13% and 11%; Fig. 5f, Extended Data Fig. 5c-d). To determine what signals are transmitted from the thalamic population to cortex, we modeled the relationship between these areas using reduced rank regression^47^, which extracts a low-dimensional channel, or communication subspace, through which thalamocortical signals flow. The largest signals within the communication subspace did not resemble the largest, rhythmic PCs in thalamus, but were instead brief transients rising immediately before obstacle traversal (Fig. 5g). Correspondingly, the thalamic factors most closely aligned with the communication subspace were not the large, sinusoidal PCs (1-2), but the smaller, transient PCs (3-4) (Fig. 5h; c.f. Fig. 5f). By contrast, thalamic factors which oscillated at twice the step frequency (PCs 7-8, Fig. 5f) were not communicated to cortex, but resided within private dimensions confined to the thalamus. These results suggest that the communication subspace amplifies transient dynamics relative to the ongoing oscillations.

**FIGURE 5.**
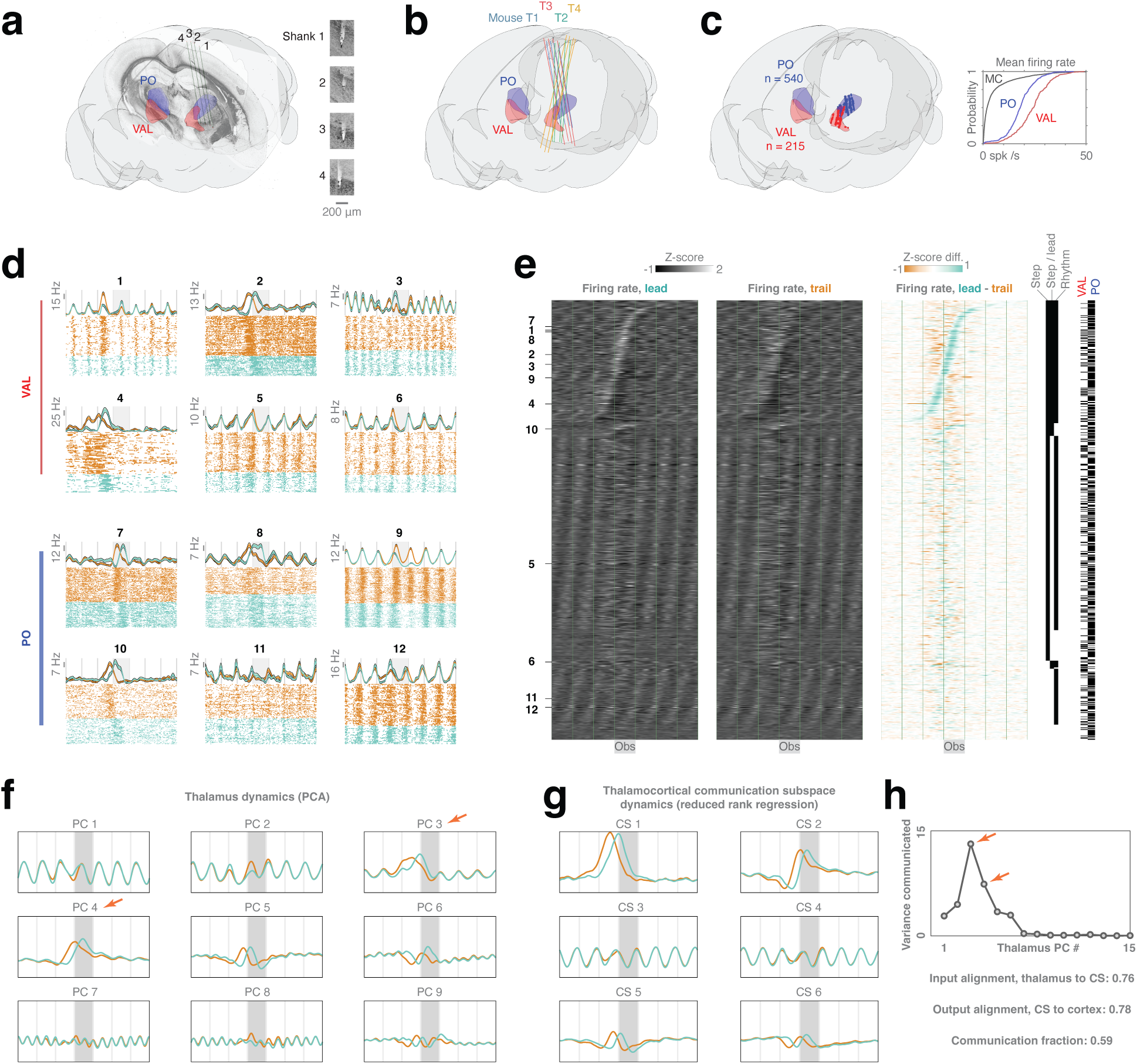
A communication subspace for thalamocortical interaction. **a**, Volumetric rendering showing the estimated position of an implanted Neuropixels 2.0 probe in thalamus, along with a coronal histological section. Insets show the tips of the four probe shanks. **b**, Probe configurations for each animal (n = 4 mice). **c**, Neuron locations in the ventral anterior lateral nucleus (VAL; n = 215 neurons, n = 8 sessions, n = 2 mice) and posterior complex (PO; n = 540 neurons, n = 19 sessions, n = 4 mice). Inset: distribution of mean firing rates in motor cortex, VAL and PO. **d**, Firing rates and raster plots for representative units in VAL and PO during obstacle traversal. Error bars denote standard error of the mean. **e**, Responses of all thalamic units (VAL and PO pooled; n = 755 neurons, n = 4985 sequences, n = 22 sessions, n = 4 mice). Conventions are the same as in Fig. 2c. **f**, Thalamic population dynamics. Each panel displays activity along one of the first nine principal components. **g**, Activity within the thalamocortical communication subspace. Each panel shows activity along one of the first six input axes. **h**, Variance communicated to motor cortex as a function of thalamic principal component number.

In our task, motor cortex must transform inputs conveying obstacle proximity and the phase of the spinal CPG into motor commands for gait modification. We have identified the representations motor cortex uses to implement this computation: a large preparatory factor synchronized with obstacle proximity, rhythmic factors entrained to locomotion both on a flat surface and during obstacle clearance, and small readout factors with the properties of limb-dependent commands. What algorithm might the cortical network use to transform the rhythmic and transient factors into a motor command? We propose a simple model that accomplishes this transformation through phase-dependent gating (Fig. 6a; Supplemental Video 3). The model consists of a system of coupled differential equations (see Methods). Dynamics within the rhythmic dimensions generate stable, step-locked oscillations, while the preparatory factor is modeled as a function of time, with a peak at the onset of contralateral swing for the step over the obstacle in the lead condition, and 0.5 cycles earlier in the trail condition (Fig. 6b, top three rows). The dynamics of the readout factor are determined by a nonlinear interaction between the rhythmic and preparatory factors, such that a transient readout signal is produced only when the preparatory input arrives early in the locomotor cycle (Fig. 6b, fourth row). Drive to the motor pools is then obtained by summing the rhythmic CPG output and this cortical readout signal (Fig. 6b, fifth row), and the resulting sum determines muscle activation (Fig. 6b, bottom row). Thus, the model transforms rhythmic dynamics, which contain no information about the obstacle, and a limb-independent input signaling obstacle proximity into a limb-dependent command which sculpts neuromotor output at the appropriate time.

**FIGURE 6.**
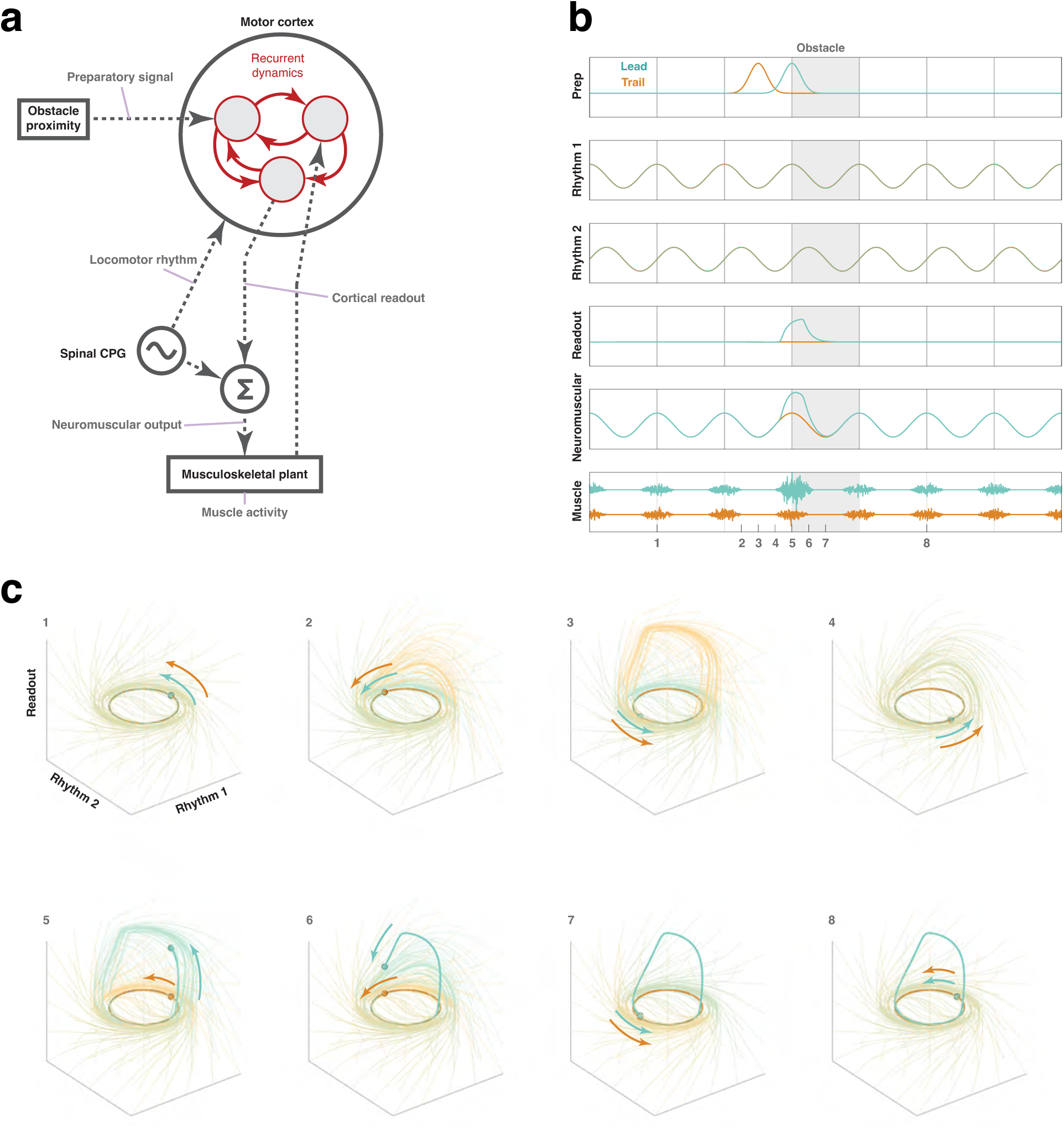
A model of cortical computation by phase-dependent gating. **a**, Block diagram illustrating the flow of information between elements of the model. **b**, Time series of model factors in lead and trail conditions. **c**, Time-dependent changes in the flow of the model vector field induced by changes in the preparatory factor. Bold lines show the trajectories of the system in the two rhythmic factors and the readout factor. Light lines show the flow computed by temporally integrating the vector field from a grid of initial conditions for the lead (blue) and trail (orange) conditions.

The phase-dependent gating mechanism underlying this transformation is illustrated in Fig. 6c, which displays the flow of the system in the rhythmic and readout dimensions at eight time points across the step sequence. During locomotion on a flat surface, the neural trajectories oscillate, and are closely matched in the lead and trail conditions (Fig. 6c, subplot 1). When the contralateral limb trails, the preparatory factor alters the vector field in the middle of the step cycle preceding the obstacle (Fig. 6c, subplots 2-4). Nevertheless, the flow near the neural state at this step phase is minimally affected by the preparatory input, and the readout factor remains nearly constant. In the lead condition, by contrast, the preparatory input reshapes the flow in precisely the same way, but does so half a cycle earlier (Fig. 6c, subplots 5-8). Consequently, the trajectory is “swept up” along the vertical axis, and a transient command for gait modification is generated at swing onset.

## 3. Discussion

Motor cortical dynamics in our task were dominated by factors that did not resemble motor commands, and are unlikely to directly control muscle activity. Instead, these factors are internal, output-null representations^21,48,49^ the cortical network uses to compute the appropriate - and much smaller - descending commands. The largest factor represented motor preparation, and was tightly locked to the approaching obstacle, reminiscent of single-neuron activity patterns reported in cat premotor areas^18,31,32^. This factor was synchronized with movement of the first forelimb to traverse the obstacle, and its amplitude did not differ between trials in which the contralateral and ipsilateral limb led. The limb independence of preparatory activity in the locomotion task contrasts with the finding that cortical activity related to the ipsilateral and contralateral limbs lies in distinct subspaces when macaques perform isolated upper limb movements^50–52^. We propose that when cortical commands must be coordinated with a subcortical CPG, as in skilled locomotion, the decision to lead with the right or left forelimb is computed within the cortical network^17^, which must therefore contain a limb-independent representation of obstacle proximity.

The strong rhythmic modulation we observed resembles the periodic patterns found in primate motor cortex during voluntary arm cycling^39,40^. The similarity in the form of the patterns, however, conceals two essential differences in their function and origin. First, while rhythmic activity in voluntary cycling is necessary for the generation of corticospinal commands, basic locomotor behavior persists after cortical dynamics are suppressed by pharmacological or optogenetic methods, or abolished entirely by lesions^13–16^, as demonstrated here. Second, in arm cycling, interactions within the cortical network generate smooth, periodic trajectories. These trajectories exhibit low tangling: the future neural states can be accurately predicted from the current state without additional information, so trajectories do not cross. Because autonomous dynamical systems (i.e., those without time-varying inputs) generate trajectories with low tangling, the observation of trajectories with this property has been regarded as evidence that the underlying neural system is minimally influenced by inputs^10,39,40,44^. During locomotion on a flat surface, motor cortical trajectories are smooth, periodic, and untangled^21^. Nevertheless, the locus of pattern generation is spinal, and the rhythm is imposed on motor cortex by periodic inputs from the spinal CPG. Thus, motor cortex can produce untangled trajectories even when operating in an input-driven, periodically-forced regime.

Our results identify a low-dimensional communication subspace through which the thalamus can drive motor cortical dynamics. Notably, this subspace is not closely aligned with the dominant thalamic dimensions, which carry step-locked rhythmic activity. Instead, it is best aligned with thalamic dimensions carrying limb-independent transients immediately in advance of the obstacle (compare Fig. 5h and 5f, PCs 3-4). Thus, in effect, the communication subspace amplifies these transients relative to ongoing rhythmic activity, enabling the generation of preparatory dynamics which prevail downstream in motor cortex (Fig. 2d, e; Extended Data Fig. 3a-c). Why does this amplification occur, and why is the preparatory factor in motor cortex so large? One possible explanation is that the preparatory factor “energizes” the cortical network, allowing it to produce rapid changes in firing rates as it computes the appropriate motor commands. In this view, the preparatory factor resembles the large, condition-independent signal that triggers movement onset in macaque motor and premotor cortex^53^. It differs, however, in that its dynamics are independent not of movement direction, but of which effector (ipsilateral or contralateral forelimb) is used to traverse the obstacle first. Furthermore, we observed that preparatory activity is maintained when the animal pauses at the barrier (Fig. 3c, d). Thus, in our task, motor preparation and initiation may not be implemented by distinct neural factors; instead, gait modification can be triggered by the coincidence of specific states along preparatory and rhythmic dimensions.

In our model, commands for gait modification are generated within motor cortex through an interaction between a limb-independent preparatory transient and ongoing rhythmic activity: different commands are produced when the transient arrives at different locomotor phases. It is possible, however, that similar phase-dependent gating mechanisms are also implemented in other CNS regions. Indeed, microstimulation of motor and premotor cortex induces different muscle responses when applied at different phases of the step cycle^54,55^, suggesting additional gating may occur at the segmental level. More broadly, we note that our model for neural control of gait modification is far from complete. For example, it is likely that posterior parietal cortex monitors the location of distant objects relative to the body by integrating visual and somatosensory inputs^17,56–59^, while the basal ganglia control task-specific motor sequences^60–62^ and steering^63^ and the cerebellum corrects for mechanical disturbances^64–66^. These nested feedback loops do not act in isolation, but interact at numerous points across the neuraxis. Furthermore, the engagement of these loops depends strongly on the task and context - whether the behavior is voluntary or automatic, precise or variable, learned or innate, and goal-directed or habitual^22,23,38^.

In the context of voluntary movement, muscle activity is often modeled as a function of motor cortical firing rates. This function, however, when estimated from recordings of cortical and muscle activity in a single task, may not fully capture the transformation of cortical dynamics into motor output^38,67^. In primates, the relationship between cortical and muscle activity can change across tasks^68,69^, even for corticospinal cells that project onto motoneurons monosynaptically^70^. Our results reveal an additional challenge in modeling cortical control of motor output. In tasks like skilled locomotion, multiple independent centers generate commands that ultimately converge on the same motoneurons. In such cases, muscle activity depends not only on the state of the cortical network, but also on the state of the spinal CPG. We propose that generalizing models of cortical control to capture interactions with subcortical circuits will enable them to explain not only discrete movements like reaching, but also a wider range of tasks which integrate automatic and voluntary processes.

## 4. Methods

### 4.1. Experimental animals

All experiments and procedures were approved by the Institutional Animal Care and Use Committee at Case Western Reserve University, and in accordance with NIH guidelines. For neural recording experiments in motor cortex, three animals were used (n=3 mice, 2 male and 1 female). These animals expressed Channelrhodopsin-2 (ChR2) in Parvalbumin (PV) neurons, and were the offspring from a cross between PV-Cre (JAX #017320) and Ai32(RCL-ChR2(H134R)/EYFP) mice (JAX #024109). For neural recordings in thalamus, four animals were used (n = 1 male mouse, PV-Cre x Ai32; and n = 3 male mice, C57BL/6, JAX #000664). For lesion and sham experiments, five animals were used in each cohort (n = 10 male mice, C57BL/6, JAX #000664). Finally, six animals were used for muscle recording experiments (n = 1 male mouse, offspring from a cross between L7-Cre2, JAX #004146, and Ai32(RCL-ChR2(H134R)/EYFP), JAX #024109; n = 3 male mice, VGAT-ChR2-EYFP line-8, JAX #014548; and n = 2 male mice, C57BL/6, JAX #000664). At the time of surgical implantation, all mice were 13–26 weeks old and weighed 22–33 g. After surgery, mice were in recovery for at least two days and closely monitored. Recordings were performed twice per day for up to three months. Animals were individually housed under a 12-h light-dark cycle at 18–24 °C and 40–60% humidity.

### 4.2. Chronic implant surgery

The following procedures were followed for all survival surgeries. Anesthesia was induced with isoflurane (1–5%, Kent Scientific), eye lubricant was applied, fur on top of the head and posterior neck was shaved, and the mouse was positioned in a stereotaxic apparatus (model 1900, KOPF instruments) on top of a heating pad. Under sterile technique, the top of the head was cleansed, lidocaine (10 mg/kg) was injected under the skin on the top of the skull, the skin was removed, the periosteum on top of the skull removed, and a custom-designed 3D-printed head post was attached with UV-cured dental cement (Re-lyX Unicem 2, 3M). Once the head post was secured, the surgery proceeded differently for neural probe implants, EMG implants, and cortical lesions. After surgery, the minimum recovery period was two days. Carprofen was administered for pain relief once per day, and the investigators monitored animal behavior, body mass, and food and water intake on a daily basis. The recovery period was extended an additional one to three days for some animals as necessary. At the time of euthanasia, animals were deeply anesthetized and transcardially perfused with 4% paraformaldehyde. For cortical recordings, a single 4-shank Neuropixels 2.0 probe was implanted in the left motor cortex of each animal^27^. The electrode was grounded with a gold pin soldered to a stainless steel wire placed in the right visual cortex. A craniotomy (dimensions 1 × 2 mm) performed using a dental drill enabled access to the forelimb area of the left motor cortex, and was centered at bregma +0.5 mm, lateral 1.7 mm. For deep brain recordings targeting the ventral anterior lateral nucleus (VAL) and posterior nucleus (PO) of the thalamus, probe insertion coordinates were calculated using the Neuropixels Trajectory Explorer (https://github.com/petersaj/neuropixels_trajectory_explorer), and a single 4-shank Neuropixels 2.0 was implanted in the left thalamus. When inserted at an angle, the probe was mounted in the Kopf Model 1950 Off-Plane Insertion Tool. The electrode was referenced with a gold pin soldered to a stainless steel wire placed in the right prefrontal cortex. A craniotomy (dimensions 1 × 2 mm) using a dental drill, was centered over the probe insertion point for each animal. See Extended Data Fig. 4 for detailed information on probe localization. Implant coordinates for thalamic recordings in each mouse are listed below.

Mouse T1 Insertion point of the most medial shank: −2.06 AP, −1.58 ML, depth 4.175 mm along the shank Angle: 0° azimuth, 90° elevation, 180° rotation

Mouse T2 Insertion point of the most lateral shank: −2.06 AP, −2.06 ML, depth 5.0 mm along the shank Angle: 41° azimuth, 75° elevation, 230° rotation

Mouse T3 Insertion point of the most medial shank: −1.10 AP, 0.70 ML, depth 5.0 mm along the shank Angle: 0° azimuth, 90° elevation, 180° rotation

Mouse T4 Insertion point of the most lateral shank: −2.37 AP, −2.40 ML, depth 5.0 mm along the shank Angle: 20° azimuth, 69° elevation, 200° rotation

The probe was implanted after the head post and ground were secured to the skull. Care was taken to leave the dura intact, and cold saline was applied continuously to reduce swelling. The probe tip was slowly inserted to a depth of 4-5 mm, silicone sealant was applied (Kwik-Sil, World Precision Instruments), and the probe assembly was secured to the head post and skull using dental cement (RelyX Unicem 2, 3M).

### 4.3. Cortical lesion

Lesions were stereotactically targeted to the left forelimb area of motor cortex. After the head post was attached to the skull, a craniotomy was made over the left hemisphere of cortex (length 2.6 mm rostral-caudal, width 2.3 mm medio-lateral; bottom right corner offset from bregma was 0.5 mm lateral and 0.4 mm caudal). Then, cortical tissue was aspirated with a sterile blunted needle (26 gauge, cut to 1.5 mm length). Bleeding was managed using cold saline, and the cavity was filled with a sterile gelatin sponge (Surgifoam, Ethicon) and sealed with silicone and then dental cement. Animals in the sham control group were not subjected to the lesion, but received a craniotomy. After euthanasia and fixation, brains were extracted and embedded in 4% agarose gel, and coronal sections of 100 µm (Vibratome, Leica VT1000 S) were imaged at 4× magnification (Slideview VS200, Evident).

### 4.4. Electromyography

Animals were implanted with Myomatrix arrays (model number RF-4×8-BVS-5) or handmade fine wire electrodes, as described previously^21^. Muscles targeted were the biceps and triceps brachii, which flex and extend the elbow, respectively, and the anterior aspect of the deltoid, which flexes the shoulder. Recordings were amplified and bandpass filtered (0.01–10 kHz) using a differential amplifier and digitized (Intan RHD2216, 16-bit, 16 channel bipolar input recording headstage), and acquired at 30 kHz (Open Ephys acquisition board and software). At the conclusion of experiments for each animal, the targeted muscles were verified post-euthanasia by dissection. For subsequent analysis of step-aligned muscle activity, the gross EMG was high-pass filtered (200–250 Hz cutoff), rectified, and convolved with a Gaussian kernel (σ = 25 ms). The EMG signal was then normalized, step-locked, processed into obstacle sequences, and subsequently averaged across trials (see the ‘Pose estimation’ subsection). For normalization, the local maxima of EMG peaks were identified with the amplitude of each peak required to be larger than its two neighboring peaks. Then, the EMG time series was divided by the median amplitude, calculated from all peaks exceeding the 90th percentile.

### 4.5. Behavioral task

Mice were placed on a custom-built motor-driven treadmill (46 cm long by 8 cm wide) that was adjusted to a constant speed of approximately 20 cm/s, as described previously^21^. Belt speed was adjusted in each session to enable smooth performance over the duration of the experiment. The treadmill apparatus was enclosed in transparent acrylic, and belt speed was monitored by a rotary encoder. Locomotion was motivated through negative reinforcement with air puffs triggered by an infrared break beam at the back of the treadmill belt. On the treadmill belt, two obstacles were taped to the belt, with an inter-obstacle distance of approximately 23 cm. The obstacles were cut-outs from standard plastic weigh boats and were approximately 6.5 cm wide and 1 cm in height. Animals were acclimated to the task over several days, until they consistently traversed the obstacles with minimal hesitation.

### 4.6. Videography and pose estimation

Images of the animal were acquired with four synchronized high-speed cameras (Black-fly, model BFS-U3-16S2C-CS, Teledyne FLIR; Vari-Focal IP/CCTV lens, model 12VM412ASIR, Tamron) at 150 Hz, as described previously^21^, 3D pose estimates were then transformed into a treadmill-belt-centered coordinate frame. Step cycles were identified by detecting threshold crossings of the forward finger velocity and upward finger position, and filtered based on step duration (60–400 ms). We then detected sequences of seven consecutive steps, where the fourth step traversed the obstacle. Sequences in which the first limb to traverse the obstacle was contralateral to the neural probe were assigned to the lead condition. Sequences in which the ipsilateral limb crossed the obstacle first were assigned to the trail condition.

### 4.7. Neural recording

During each session, the headstage was connected to the probe and secured to the headpost. For each animal, we surveyed neural activity across all shanks and channels to determine the optimal channel recording configurations for the target region. During behavior, cortical recordings were acquired from four banks of 96 channels spanning all shanks, at depths ranging from 0.41-1.27 mm from the surface of the brain. For thalamus, recordings from separate animals were similarly acquired in four banks of 96 channels at relatively greater depths (see Extended Data Fig. 4) than cortex. Recordings were amplified, bandpass filtered (0.1-10 kHz) and acquired at 30 kHz using National Instruments PXIe/PCIe-8281 controller module and Open-Ephys system (3rd generation acquisition board and GUI software).

### 4.8. Spike sorting

Single units were identified using Kilosort 4.0^71^ and manually curated with the Phy 2.0 GUI (https://github.com/cortex-lab/phy). Single units were isolated based on spike waveforms, the presence of refractory periods greater than 1 ms, the stability of spike amplitude over the session, and isolation of the cluster in feature space. Spike time cross-correlation was used to remove duplicated neurons. In some experiments, single neurons were tracked across multiple sessions (Fig 3g-h; Fig 4). Binary files were concatenated and timestamps were adjusted accordingly. To minimize the effects of probe drift, concatenated sessions were often recorded during the same day, and up to 2 days apart. Single units identified by Kilosort were required to have continuous spiking activity across sessions and were otherwise excluded.

### 4.9. Optogenetic perturbation of barrel cortex

In PV-Cre x Ai32 mice, photostimulation of paravalbumin-expressing interneurons induces local inhibition in barrel cortex^69^. Optical fibers (FT200UMT, core diameter 200 µm, ThorLabs) were glued inside ceramic ferrules (catalog number CFLC230-10, ThorLabs) and positioned onto the skull over a thin layer of transparent dental cement (Optibond, Kerr), enabling optical access to the brain. The optical fiber was chronically implanted during the neural implant surgery over the left barrel cortex (bregma −1.2 mm, lateral 3 mm)^16,72^. Optogenetic perturbation with a 473 nm wavelength laser (Opto Engine LLC) was delivered with sinusoidal waves at 40 Hz for 1 second, with a random 3-4 second inter-stimulus interval (Fig. 3e-f). The laser was triggered by an external signal generator controlled with custom LabVIEW software. Power measured at the tip of the optical fiber was calibrated before the surgical implantation. Laser power was fixed at a constant value within each session. Across sessions, the power ranged from 2.5 to 10 mW.

### 4.10. Behavioral manipulations

To remove whisker sensory input during obstacle traversal, the whiskers were trimmed in two separate stages (Fig. 3g). After a standard behavior session with intact whiskers, the animal was anesthetized under isoflurane and placed on a heating pad, then the whiskers were completely trimmed on the right side, contralateral to the motor cortical probe. We then performed behavioral experiments after five hours of rest, and again the following morning. Next, the remaining whiskers on the left side of the snout and under the jaw were trimmed. The animal was then tested in two additional sessions. To remove visual input during obstacle traversal, we tested the animal in complete darkness (Fig 3h). After a standard behavior session with the room lights on, all visible light during the next session was removed. The treadmill and recording rig were completely enclosed with sheets of opaque high density polyethylene and the room lights were turned off. Each session was between 25-30 minutes in duration, and occurred after the whiskers had regrown. Animals were unable to perform the task in the dark while the whiskers were trimmed.

### 4.11. Passive limb movement

After a standard behavioral session on the treadmill, the experimenter handled the mouse and applied passive forelimb movements (Fig 4a). The mouse remained in the investigator’s left hand, and once the mouse became calm and encouraged with juice reward, a manipulandum (cotton swab) was placed under the right paw and a series of passive forelimb movements were applied as the animal remained at rest. The paw was moved in forward cycling patterns. The behavior was recorded using a single camera from the treadmill rig (modified ROI and frame rate, 468 x 466 pixels and 60 Hz, respectively). In DeepLabCut, the positions of 8 landmarks were tracked, corresponding to the nose, eye, ear, elbow, wrist, and fingertip on the right side, the cotton swab head and cotton-wood junction. In total, 854 labelled frames were used to train the model for passive limb movement. To identify forward movement cycles, upward velocity thresholding was used, and each cycle started at zero-velocity crossing. Cycles were then filtered based on distance of the fingertip to the cotton swab, and low movement variability of the nose, eye and ear. Additionally, cross-correlation between finger tip upward and forward velocity was used to select forward cycling trials. Simultaneously, electrophysiological recordings in the motor cortex using the same channel configurations as during locomotion were performed.

### 4.12. Probe alignment and identification of thalamic nuclei

After fixation, the brain was transferred to 30% sucrose in a phosphate buffer at 4°C for 36–48 hours until the tissue dehydrated and sank. Next, brains were embedded in mounting medium (OCT compound, Tissue-Tek, Sakura) and frozen to −80 C using dry ice. Coronal cryosections were cut at 25 µm thickness (Leica CM3050S), rinsed with phosphate buffer, and mounted with antifade medium (Vectashield, Vector Laboratories). Mounted sections were imaged at 4x magnification using a slide scanner (Slideview VS200, Evident). Slide-scanned images were registered to the Allen Mouse Brain CCFv3 (2017 version; 25 µm isotropic resolution) using QuickNII and VisuAlign based upon correspondence of anatomical landmarks. Tracks cut by each electrode shank, as well as the shank tips, were visible in tissue sections, and the track locations were annotated manually. All annotated track points and endpoints were mapped into CCFv3 coordinates using the previously-computed registration. A 3D line was then fit through the points for each shank using a least-squares fit. The endpoints of the fitted line were defined as the projections of the most dorsal and most ventral annotated points onto the fitted direction.

Because tissue deformation introduced differences between the section scale and the Allen CCF reference volume, a stretch ratio was calculated for each mouse to correct the electrode spacing in CCF space. The stretch ratio was defined as the mean brain image aspect ratio before the stretch divided by the atlas aspect ratio across all sections within the probe span. This ratio was applied multiplicatively to the known pitch between the electrodes (15 µm) and to the tip-to-first-electrode offset (175 µm) to yield the stretched spatial pitch of the electrode in CCF space. Electrode positions were calculated along each shank trajectory, with the most ventral electrode near the tip calculated first. At each row, two electrode coordinates were calculated by applying medial-lateral offsets of −5 µm and +27 µm that are perpendicular to the shank axis to match the probe geometry. Each electrode’s CCF coordinates were stored and used to query the Allen CCF-annotated volume to retrieve the corresponding brain structure labels. The reconstructed electrode coordinates and their histological labels were validated against an electrophysiological survey of activity across each shank. During the earlier in-vivo recording phase, we sequentially recorded 2-5 minutes of data from each bank of channels in the brain as the mouse stood on a slowly-moving treadmill belt. This enabled us to build a functional map of spiking activity across the length of each shank. After sacrificing the animal and performing the analysis of histological sections described above, a small offset was therefore applied to the estimated location of each shank tip to account for spatial discrepancies (0 to 270 µm adjustment). After fine-tuning, these region assignments were used to categorize recording channels and to assign sorted neurons to specific anatomical locations.

Data analysis

### 4.13. Firing rate modulation for single neurons

Firing rates over the full experimental session were computed using Gaussian smoothing (σ = 25 ms). Using step sequence segmentation from kinematic data (described above), smoothed firing rate curves were extracted for each step using linear interpolation between the start of swing and end of stance. Sequence-aligned averaged firing rates were computed across the seven identified steps to create peri-event time histograms (Fig. 2b). For comparisons across neurons, firing rates were Z-scored based on activity from the identified step sequences (Fig. 2c, Extended Data Fig. 2a-c).

Firing rates on individual strides were computed by dividing the number of spikes in the cycle by the cycle duration. To determine how firing rates were influenced by step number within the sequence, the lead vs. trail condition, and the interaction between these two factors, we performed a two-way ANOVA for each neuron. Because the animal’s behavior on the initial and final steps of the sequence was nearly identical in the lead and trail conditions, we rarely observed a significant main effect of lead vs. trail (3% of recorded neurons). Entrainment to the locomotor rhythm was determined using a Rayleigh test for each cell. The Benjamini-Hochberg correction for multiple comparisons was applied to the p-values obtained from the ANOVAs and Rayleigh tests.

For passive limb movement, the cycling frequency was slower than the stepping frequency during locomotion (range 0.5-3 Hz, as compared to 3-5 Hz, respectively). Therefore, during passive movements, firing rates were computed over the session with a longer Gaussian smoothing kernel (σ = 50 ms). Rhythmicity was calculated using the Rayleigh test, and the tuning strength and preferred phase were defined as the mean resultant length and angle, respectively. The Benjamini-Hochberg correction was then applied.

### 4.14. Neural factors in motor cortex

For each experimental session, principal component analysis (PCA) was performed on the Z-scored, trial-averaged firing rates for all simultaneously-recorded neurons. Z-scored firing rates on single trials were then projected onto the coefficients. For subsequent analyses, we identified three types of population-level factors, each of which was a linear combination of firing rates. The weights for each factor were scaled to have a norm of unity.

(1) Preparatory factor, defined to be the first principal component of cortical activity.
(2) Two rhythmic factors, obtained by linear regression of the cosine and sine of step phase on the concatenated single-trial scores for the first 20 cortical principal components.
(3) Three readout factors, obtained by linear regression of the residual EMG template for each muscle on the first 20 cortical principal component scores. The residual EMG template was obtained by subtracting the average step-locked EMG signal for steps on a flat surface from the step-locked signal within a two-stride window centered on the middle of the cycle traversing the obstacle. This template was averaged over all datasets for each muscle (Extended Data Fig. 1c).

### 4.15. Comparison of responses to active and passive movement

Single neurons recorded during locomotion on a flat surface and passive movement of the contralateral forelimb were linked (see Spike sorting above), and activity was compared between conditions. For active movement, we selected flat steps cycles and excluded those preceding and crossing the obstacle. Tuning strength and preferred phase were defined as the mean resultant length and angle for the active or passive movement phases sampled at spike times (Fig. 4b). Within each condition (active and passive), we computed the correlations between the firing rates of all pairs of neurons. We then obtained condition-specific orderings of neurons using agglomerative hierarchical clustering (Euclidean distance and complete linkage) on the correlation matrix (Fig. 4d). Active and passive subspaces were obtained by linear decoding of the cosine and sine of the locomotor rhythm and the passive movement rhythm, respectively (Fig. 4e-f).

### 4.16. Modeling thalamocortical communication

Analysis of single units and principal components for the thalamus data proceeded as for the cortical data. To capture the relationship between experimentally-observed thalamic inputs and cortical activity, we fit a reduced rank regression model of the form:

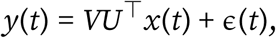

where y(t) represents the activity of motor cortical neurons, x(t) represents thalamic neurons, U and V are rank r matrices, and ɛ(*t*) is an error term. In this model, thalamic inputs pass through a low-dimensional bottleneck, or communication subspace, through which they influence cortical dynamics. We fit and analyzed this model using the procedures and code described in Wu and Pillow^47^ (arXiv, 2025; https://github.com/bichanw/RRR). Briefly, we fit U and V using a ridge estimator with ridge penalty λ = 10e3, and r = 6. Activity within the communication subspace was obtained by projecting y(t) onto the columns of U (Fig. 5g). The alignment between thalamic principal components and the communication subspace (i.e., the variance communicated; Fig. 5h) was defined as:

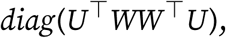

with *W = UV⊤*.

Finally, the overall alignment between the communication subspace and the major activity modes within the input (thalamic) and output (cortical) areas was calculated using the normalization procedures described in Wu and Pillow^47^.

### 4.17. Computational model

We modeled the transformation of oscillatory dynamics and a limb-independent transient into a limb-dependent cortical command using a system of coupled differential equations (Fig. 6). The limb-independent preparatory input was modeled as a function of time,

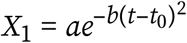

where *t* ∈ [–3, 4] represents the time in the step sequence, and a = 2.5, b = 30. The peak *t*_0_ occurred at 0 in the lead condition, and at −0.5 in the trail condition. The oscillatory dynamics were generated using

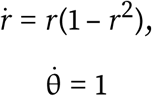

and the rhythmic factors were defined as

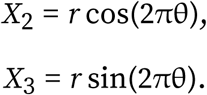

The readout factor was produced through a nonlinear interaction between the preparatory and rhythmic factors:

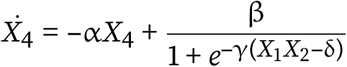

with α = 10, β = 20, γ = 10, and δ = 0.5. Initial conditions at *t* = –3 were set to *X*_2_(–3) = 1, *X*_3_(–3) = 0 and *X*_4_(–3) = –3. The equations were integrated using the Euler method with a time step of 0.002. Finally, the neuromotor drive was obtained by adding *X*_2_ and *X*_4_.

## 5. Author contributions and acknowledgements

E.A.K., K.C., and B.A.S. designed the experiments. E.A.K. performed the cortical and EMG recordings and the lesion experiments. K.C. and E.A.K. performed the thalamus recordings. E.A.K., K.C., and B.A.S. analyzed the data. K.C. developed the probe localization pipeline and performed histological procedures. E.A.K., K.C., and B.A.S. interpreted the results and wrote the paper. We thank Trevor Drew, Andrew Pruszynski, Hillel Chiel, Andrew Spence, and Torsten Ullrich for discussions, Ian Torres, Raquel Lopez de Boer, and Polyxeni Philippidou for advice on histology, and Keenan Hope for assistance with pilot experiments. This work was supported by the National Institute for Neurological Disorders and Stroke (R01NS129576, PI: Sauerbrei), Natural Sciences and Engineering Research Council of Canada (Postdoctoral Fellowship: Kirk), and the Case Western Reserve University School of Medicine. This project made use of the High Performance Computing Resource in the Core Facility for Advanced Research Computing at Case Western Reserve University. Schematic drawings of mice in Fig. 3 and 4 are adapted from images in a public repository (https://doi.org/10.5281/zenodo.3925915,https://doi.org/10.5281/zenodo.3925902). These images are available under a Creative Commons 4.0 license (CC-BY). The authors declare no competing or financial interests.

## 6. Code and data availability

On publication of the manuscript, data and code will be deposited for public access on Dryad.

---

**Extended Data Figure 1:**
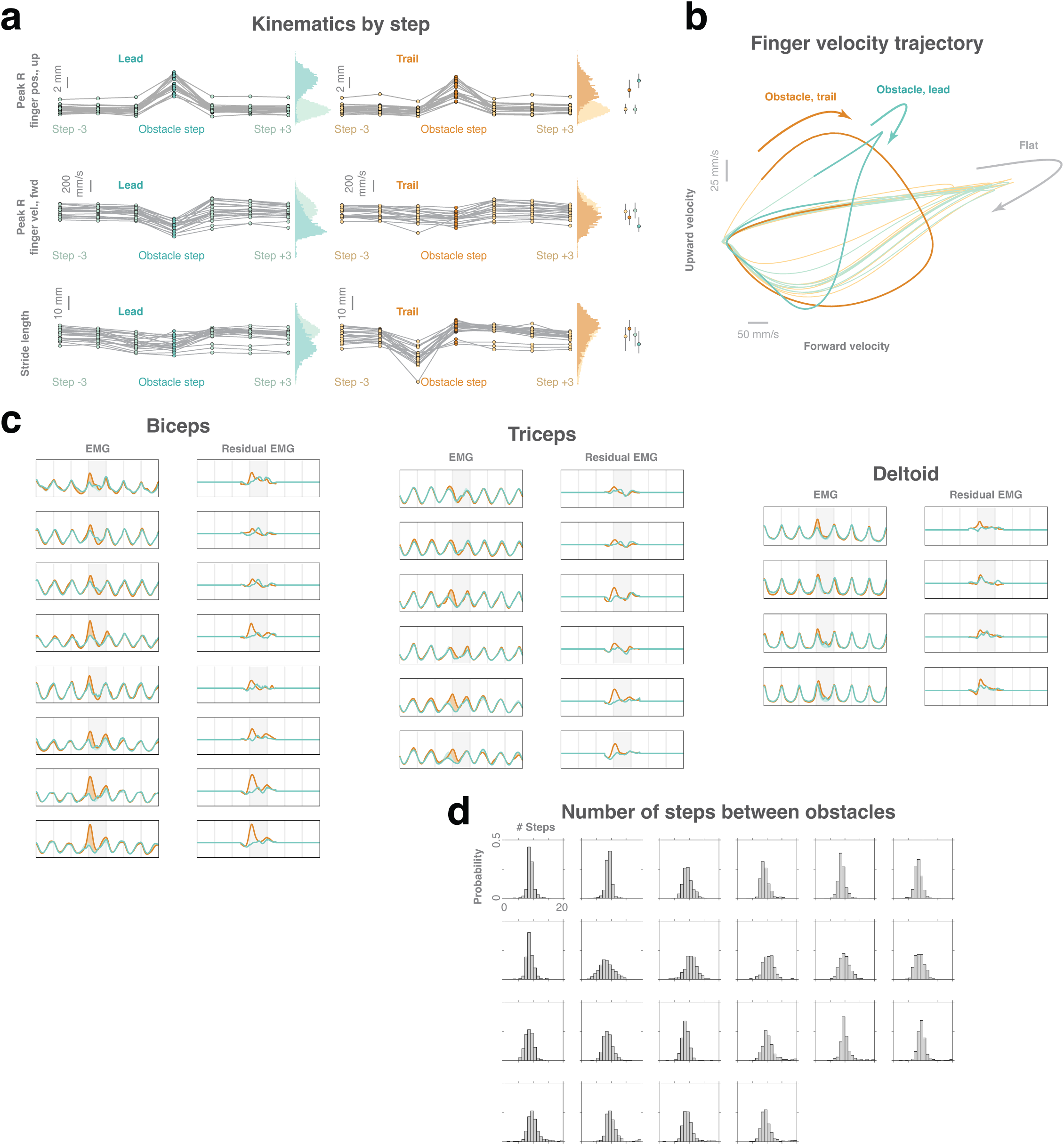
Kinematics and muscle activity during obstacle traversal. **a**, Median peak finger height (top row), finger forward velocity (middle row), and right forelimb stride length (bottom row) on each step in the sequence. Each curve shows data from one experimental session (n = 22 sessions, n = 3 mice). Histograms show the distribution of values across all single trials for flat steps (light) and obstacle traversals (bold). Inset at right shows the quartiles for each condition (lead and trail, and flat and obstacle). **b**, Velocity trajectory of the right fingertip during the step sequence. The obstacle traversal is bolded (lead and trail), and the velocity is shown along the forward and upward directions. **c**, Electromyogram (EMG) and residual EMG after subtraction of the flat-surface template. Data are plotted as the mean for lead and trail sequences, each row corresponds to a different session, and sessions are grouped by muscle (biceps brachii, n = 8 sessions, n = 3 mice; triceps brachii, n = 6 sessions, n = 3 mice; anterior aspect of the deltoid, n = 4 sessions, n = 2 mice). **d**, Histograms show the number of steps between obstacle traversals for each session (n = 22 sessions, n = 3 mice).

**Extended Data Figure 2:**
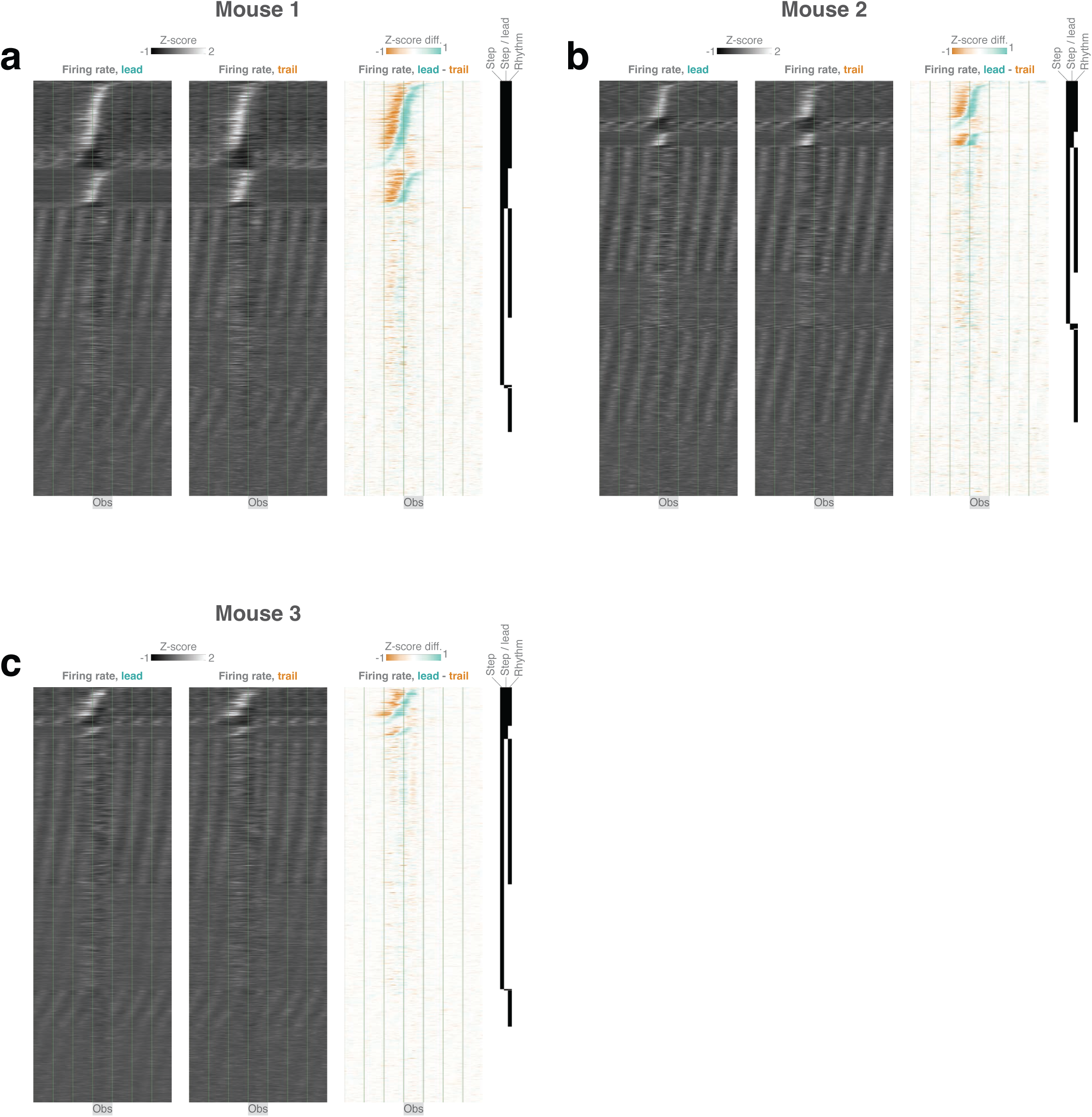
Motor cortical activity during the skilled locomotion task for each experimental animal. **a**, Task-related activity for all neurons recorded in mouse 1 (n = 1982). Conventions are as in Fig. 2c. **b**, Responses for mouse 2 (n = 1919). **c**, Responses for mouse 3 (n = 1695).

**Extended Data Figure 3:**
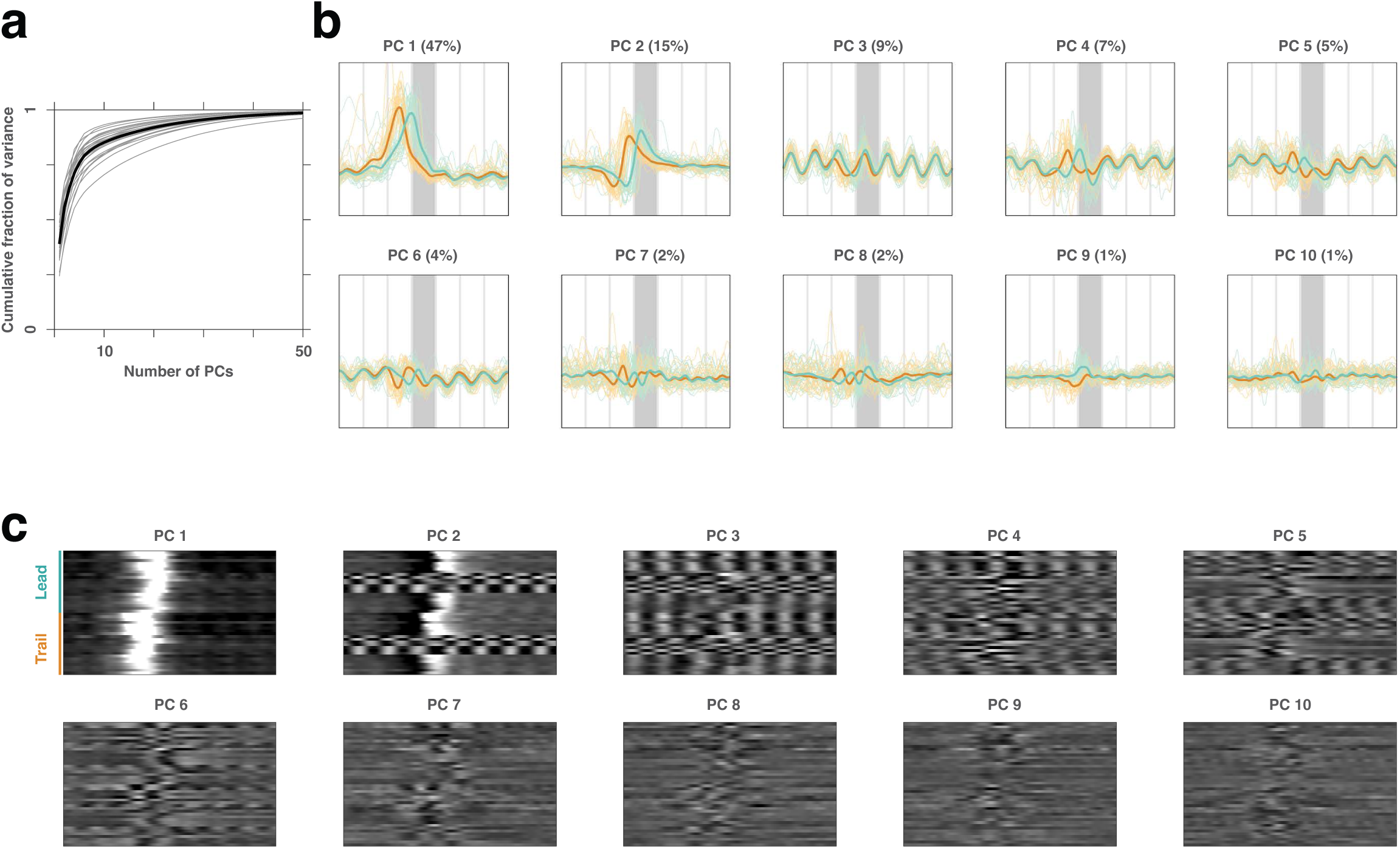
Cortical population dynamics during the skilled locomotion task in principal component coordinates. **a**, Cumulative fraction of firing rate variance explained as a function of number of components. Each gray curve corresponds to one session, and the black curve corresponds to the mean across sessions. **b**, Single-trial scores for the first ten principal components for a representative session (n = 239 neurons, n = 311 sequences, n = 1 mouse). **c**, Trial-averaged principal components for all sessions (n = 5596 neurons, n = 5519 sequences, n = 22 sessions, n = 3 mice).

**Extended Data Figure 4:**
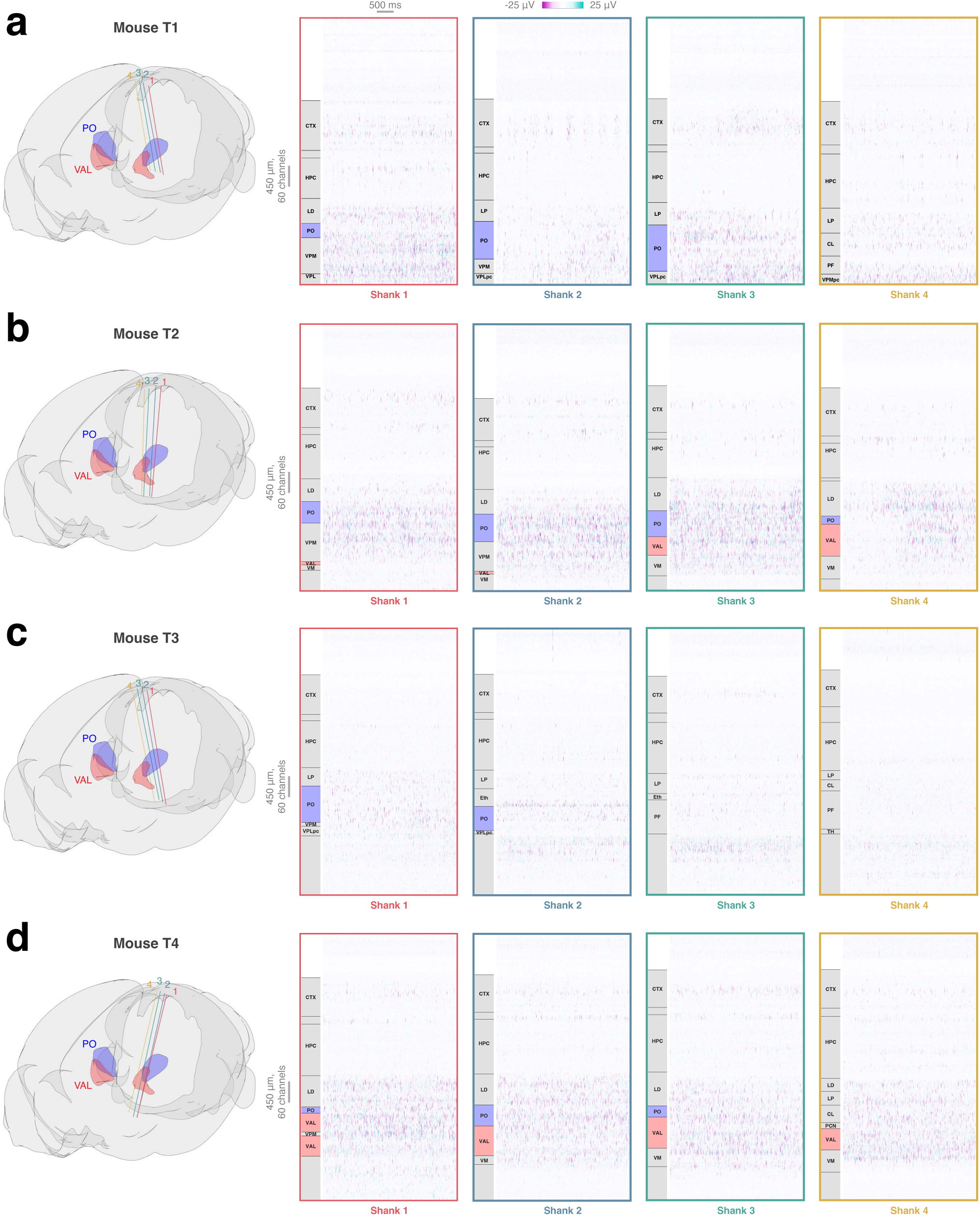
Localization of thalamic neurons. **a**, Histological and electrophysiological localization of the probe implanted in mouse T1. Left: estimated locations of the four Neuropixels 2.0 probe shanks in the Allen Mouse Brain Common Coordinate Framework. Right: raw electrophysiological data from all channels on each shank. The mouse was slowly stepping and stopping on the treadmill during this recording. Brain area annotations are shown in left insets for each panel. Units from the posterior complex (PO) were recorded in all animals (T1-T4, n = 4 mice), and units from the ventral anterior lateral nucleus (VAL) in two animals (T2 and T4). **b**, Probe localization for mouse T2. Conventions as in **a**. **c**, Probe localization for mouse T3. **d**, Probe localization for mouse T4. Abbreviations: CL, central lateral nucleus; CTX, cortex; Eth, ethmoid nucleus; HPC, hippocampal formation; LD, lateral dorsal nucleus; LP, lateral posterior nucleus; PCN, paracentral nucleus; PF, parafascicular nucleus; PO, posterior nucleus; TH, undefined thalamus; VAL, ventral anterior lateral nucleus; VM, ventral medial nucleus; VPL, ventral posterolateral nucleus; VPLpc, ventral posterolateral nucleus parvicel-lular part; VPM, ventral posteromedial nucleus.

**Extended Data Figure 5:**
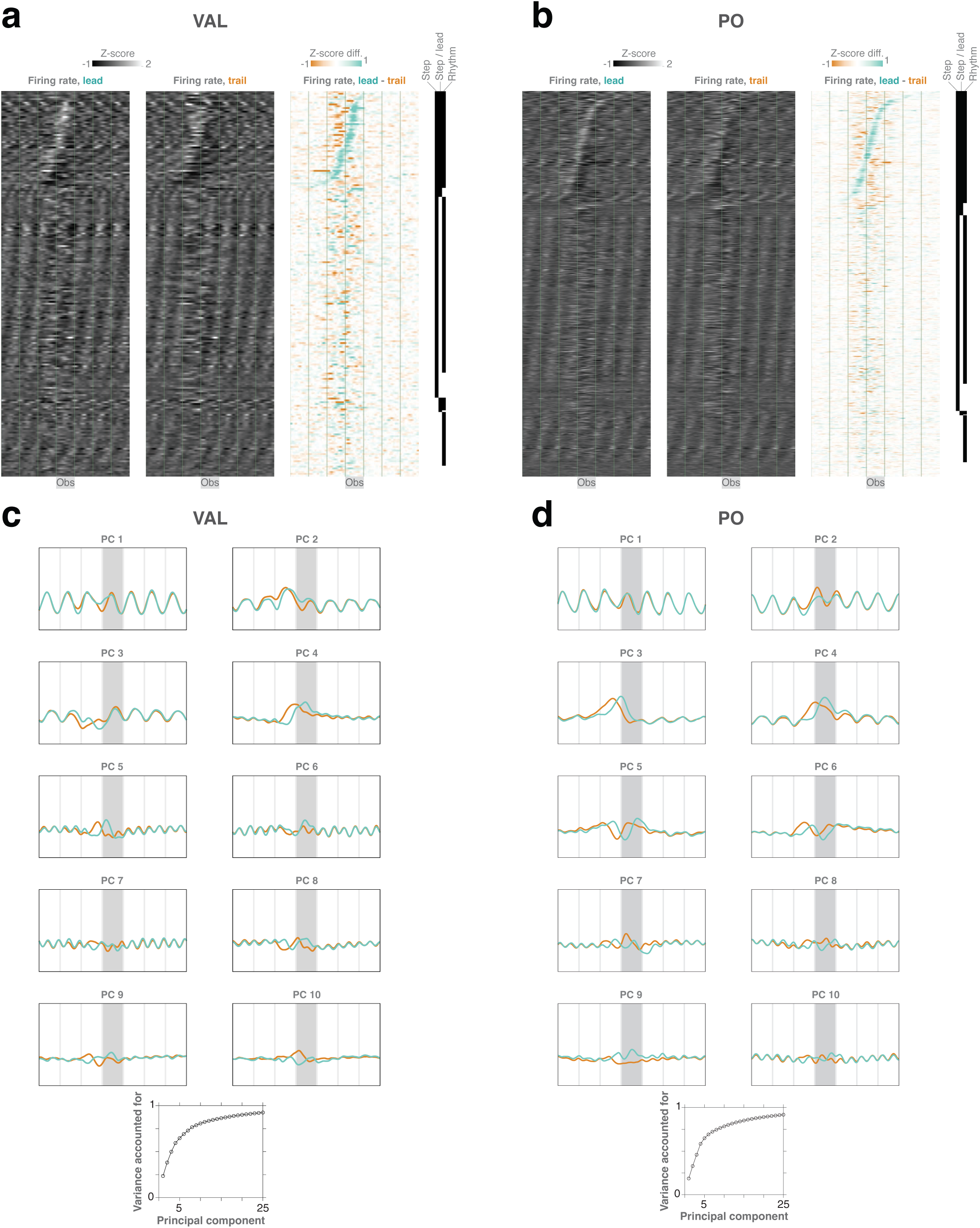
Single-unit and population dynamics in VAL and PO. **a**, Task-related activity for all neurons recorded in ventral anterior lateral nucleus (VAL, n = 215 neurons, n = 8 sessions, n = 2 mice). **b**, All neurons recorded in posterior nucleus (PO, n = 540 neurons, n = 19 sessions, n = 4 mice). Conventions are as in Fig. 2c. **c**, Scores for the first ten principal components in VAL (n = 215 neurons, from panel a). **d**, Scores for the first ten principal components in PO, and cumulative variance by PC number (n = 540 neurons, from panel b). Cumulative variance accounted for is shown in lower insets.

## Notes

### Competing Interest Statement

The authors have declared no competing interest.

### Summary of Updates

Major updates: (1) Thalamus recordings and the identification of a thalamocortical communication subspace (Fig. 5; Fig. E4-5). (2) Lesion experiments demonstrating the necessity of motor cortex for the skilled locomotion task (Fig. 1c-d). (3) Identification of cortical readout factors by decoding of residual EMG (Fig. 2d-e). (4) Additional quantification of limb kinematics and EMG (Fig. E1). (5) Some changes to the text throughout.

